# RNA-Seq analysis and annotation of a draft blueberry genome assembly identifies candidate genes involved in fruit ripening, biosynthesis of bioactive compounds, and stage-specific alternative splicing

**DOI:** 10.1101/010116

**Authors:** Vikas Gupta, April D. Estrada, Ivory Blakley, Rob Reid, Ketan Patel, Mason D. Meyer, Stig Uggerhøj Andersen, Allan F. Brown, Mary Ann Lila, Ann E. Loraine

## Abstract

**Background:** Blueberries are a rich source of antioxidants and other beneficial compounds that can protect against disease. Identifying genes involved in synthesis of bioactive compounds could enable breeding berry varieties with enhanced health benefits.

**Results:** Toward this end, we annotated a draft blueberry genome assembly using RNA-Seq data from five stages of berry fruit development and ripening. Genome-guided assembly of RNA-Seq read alignments combined with output from ab initio gene finders produced around 60,000 gene models, of which more than half were similar to proteins from other species, typically the grape *Vitis vinifera*. Comparison of gene models to the PlantCyc database of metabolic pathway enzymes identified candidate genes involved in synthesis of bioactive compounds, including bixin, an apocarotenoid with potential disease-fighting properties, and defense-related cyanogenic glycosides, which are toxic.

Cyanogenic glycoside (CG) biosynthetic enzymes were highly expressed in green fruit, and a candidate CG detoxification enzyme was up regulated during fruit ripening. Candidate genes for ethylene, anthocyanin, and 400 other biosynthetic pathways were also identified. Homology-based annotation using Blast2GO and InterPro assigned Gene Ontology terms to around 15,000 genes. RNA-Seq expression profiling showed that blueberry growth, maturation, and ripening involve dynamic gene expression changes, including coordinated up and down regulation of metabolic pathway enzymes and transcriptional regulators. Analysis of RNA-seq alignments identified developmentally regulated alternative splicing, promoter use, and 3’ end formation.

**Conclusions:** We report genome sequence, gene models, functional annotations, and RNA-Seq expression data that provide an important new resource enabling high throughput studies in blueberry. RNA-Seq data are freely available for visualization in Integrated Genome Browser, and analysis code is available from the git repository at http://bitbucket.org/lorainelab/blueberrygenome.

## Introduction

A diet rich in blueberries can help protect against diabetes [1], cardiovascular disease, and age-related cognitive decline [2, 3]. Molecular or biochemical mechanisms underlying these positive health benefits are not known, but most research has thus far focused on the antioxidant and anti-inflammatory properties of polyphenolic phytochemicals that accumulate as fruit ripen. Blueberry fruit are an especially rich source of polyphenolic anthocyanin pigments, which give blueberries their distinctive color. Of these, malvidin, delphinidin, and peonidin are the most abundant by weight [4]. The relative abundance of anthocyanin species can differ between genotypes [5, 6], and in vivo research has shown that different anthocyanins affect biological systems in different ways [7-9], suggesting that berry varieties may offer distinct health benefits. Blueberries also contain relatively large amounts of quercetin [4], another polyphenolic that may have beneficial effects in Alzheimer’s disease [10] and inflammation-related disorders [11]. Berries may also contain other as-yet undiscovered beneficial phytochemicals that could interact with anthocyanins or other compounds to potentiate biological efficacy [6, 12]. Genomic studies that catalog the full genetic repertoire of blueberry could enable greater understanding of bioactive compounds, a necessary step toward developing new varieties bred for health benefits.

Blueberries are in the Cyanococcus section of family *Ericaceae*, genus *Vaccinium*, which also includes cranberry (*V. macrocarpon*), lingonberry (*V. vitis-idaea*), and more than 400 other species [13]. Commercially harvested blueberry species in North America include lowbush (wild) blueberry *Vaccinium angustifolium*, a low, spreading shrub grown in managed stands in the Northern US and Canada, and the highbush blueberry species *Vaccinium* c*orymbosum* and *Vaccinium ashei*, which are larger shrubs grown in orderly rows in orchards and require annual pruning to maintain productivity. Of the two highbush species, *Vaccinium corybosum* is the most widely grown, while *Vaccinium ashei* is grown solely in the Southern US. *V. corymbosum* was first domesticated in the early 20th century by USDA scientist Fredrick Coville working with New Jersey farmer Elizabeth White, who recruited local pickers to locate wild berry plants with unusually large fruit. Coville’s breeding these early wild selections produced varieties suitable for commercial production, some of which are still grown today. Both lowbush and *V. corymbosum* highbush blueberries are deciduous and require a period of low temperatures during the winter season to induce flowering the following spring. To expand the range where highbush blueberries can be grown commercially, breeding programs have selected varieties with reduced chilling requirement, leading to development of sub-varieties called “southern highbush” because they require fewer days of colder temperatures to trigger flowering. Ploidy levels of berry species range from diploid to hexaploid, and most varieties of highbush berry contain genetic material introduced from diverse genotypes and species, including *V. darrowii* Camp (evergreen blueberry) and *V. arboreum* Mar. (sparkleberry), as well as rabbiteye and lowbush blueberry. Although there is a great diversity across varieties, highbush blueberry plants within the same cultivar are highly uniform, as all are clones propagated from a single selection. Thus sequences collected from individuals from the same cultivar are expected to be highly homogenous with few differences between individuals.

Estimates based on flow cytometry predict that a haploid blueberry genome is around 600 million bases, five times the size of the *Arabidopsis thaliana* genome [14]. In a related study, a draft genome assembly of a diploid northern highbush blueberry was generated using HiSeq Illumina reads [15]; the unassembled sequences are available from the Short Read Archive under accession SRA053499. This draft assembly consists of 225,479 contigs organized into 13,757 scaffolds with an N50 scaffold size of 145 kb, meaning that at least half of the sequence data is organized into scaffolds of 145 kb or larger. Plant genes are typically smaller than 2 kb, and intergenic regions are often smaller, which means that a 145 kb or larger contig could accommodate 50 or more genes. Although the genome assembly is still a work in progress, its large N50 make this draft assembly an important new resource for RNA-Seq analysis and gene discovery in blueberry.

To date, blueberry improvement efforts have focused on agronomic traits, such as ability to withstand mechanical harvesting, or consumer-focused traits, such as berry size, flavor, and mouth feel. Due to rising consumer interest in the health-protective effects of blueberries and other fruits and vegetables, breeding for nutritional and health-protective qualities may become practical in the near future. Breeding a more healthful berry will require more complete knowledge of genes encoding enzymes of secondary metabolism as well as their putative regulators. Toward this end, we performed high-throughput transcriptome sequencing (RNA-Seq) and differential gene expression analysis of five stages of berry development and ripening. Genome data, RNA-Seq expression profiles, and functional annotations have been made publicly available and will provide an important new resource for interpretation of high-throughput data from blueberry species.

## Data Description

### Berry collection and RNA extraction

Blueberry samples were collected from the field from four- or five-year old blueberry plants growing at the North Carolina Department of Agriculture Piedmont Research Station. Plants were labeled by row and position within the row; for example, plant 2-41 occupied position 41 within row 2. All samples intended for RNA extraction were flash-frozen on liquid nitrogen in the field immediately after collection and stored at -80° C until use. For RNA extraction, whole berry samples were ground to powder in a mortar and pestle with liquid nitrogen and total RNA was extracted using the Spectrum Total Plant RNA Kit (Sigma). Extracted RNAs were treated with DNase I prior to library construction using RNAase-Free DNase (catalog number 79254) from Qiagen.

### 454 library construction and sequencing for May 2009 samples

Two libraries were prepared from samples of green and ripe fruit respectively from plants of the O’Neal variety of southern highbush blueberry. The green fruit library was prepared from a mix of unripe, green fruits of varying sizes harvested on May 18, 2009 from plant 2-41. The ripe fruit library was prepared from ripe fruits harvested on June 15, 2009 from plants 2-40, 2-41, and 2-42, also of the O’Neal variety. The libraries for sequencing were constructed using the SMART PCR cDNA synthesis kit from CloneTech. The 3’ and 5’ primers used in first strand cDNA synthesis were aagcagtggtatcaacgcagagtact(30)VN and aagcagtggtatcaacgcagagtacgcggg, respectively, where V was a, g, or c and N was any nucleotide. The products of first strand cDNA synthesis were amplified using a polyA disruption PCR primer designed to introduce non-A bases in the polyA tail region of the cDNA, since homopolymeric sequences are difficult to sequence using the 454 technology. The polyA disruption primer sequence was attctagaggccgaggcggccgacatgt(4)gtct(4)gttctgt(3)ct(4)VN, where numbers in parentheses indicate the number of times the preceding base appeared in the sequence and V was a, g, or c. The sequence of the 5’ primer used for second strand cDNA synthesis was aagcagtggtatcaacgcagagt. The two libraries were sequenced in two sectors of the same plate on a 454-GS FLX Titanium sequencer (454 Life Sciences, Roche Diagnostics, USA) at the David H. Murdock Research Institute. Sequence data are available from the Short Read Archive under accession SRP039977.

### 454 library construction and sequencing for May 2010 samples

Green and ripe berries were harvested from O’Neal variety plant 2-42 on April 29, 2010 and May 26, 2010, respectively. For sequencing and library construction, samples of total RNA were sent to the North Carolina State University Genome Sciences Laboratory (GSL). Each sample was used to synthesize two libraries, which were sequenced on the same plate of a 454-GS FLX Titanium sequencer. Libraries were synthesized at the GSL following the protocol reported previously [16]. Sequence data are available from the Short Read Archive under accession SRP039977.

### Berry collection, library synthesis, and sequencing of berry development samples

Berries from five stages were selected from three plants (3-33, 2-41, and 2-42) of the O’Neal variety of southern highbush blueberry *Vaccinium corymbosum*. We designated the five stages as pads, cups, green, pink, and ripe. The “pads” and “cups” stages corresponded approximately to stages S1/S2 (pads) and S3/S4 (cups) described in [17]. Green fruit were fully rounded green berries, pink berries were partially pigmented but still firm, and ripe berries were fully colored and soft. Samples were collected during the growing season of 2011. Pads were collected on April 4, cups on April 19, mature green fruit on April 28, pink fruit on May 20, and ripe fruit on June 2. Following RNA extraction and DNAase I treatment (as described in the previous section) libraries were synthesized using the TruSeq A kit (catalog number FC-121-1001) from Illumina (Illumina, USA) following the manufacturer’s instructions. Libraries were synthesized using different TruSeq adapters to allow multiplexing, combined and then sequenced in three lanes with five libraries per lane using a HiSeq sequencer from Illumina. Sequence data are available from the Short Read Archive under accession SRP039977.

### Berry collection, library synthesis, and sequencing of berry cultivars

Fully ripe and green berries from four berry cultivars (Pamlico, Lenoir, O’Neal and Ozark Blue) were harvested in 2009 from plants growing at the Piedmont Research Station (same field as above) and frozen on liquid nitrogen in the field. RNA was extracted as described in the preceding section. Libraries were synthesized using the mRNA-Seq Sample Preparation kit (catalog number RS-930-1001) from Illumina following manufacturer instructions. Libraries were sequenced using paired-end, 76 cycle sequencing at the UNC Chapel Hill Lineberger Cancer Research Center on a GAIIx sequencer from Illumina. Sequence data are available from the Short Read Archive under accession SRP039971.

### Sequence processing and alignment

Prior to alignment, all sequences were trimmed to remove low-quality bases at the 5’ and 3’ ends of sequences. Single-end Hiseq reads (100 bp) from the berry fruit development series were trimmed to 85 bases to remove lower quality bases. Five bases were trimmed from the three prime end and 10 bases were trimmed from the five prime end of each read using the FASTX-Toolkit from Galaxy [18]. Similarly, the 76 bases long paired-end GAIIx sequences were trimmed to 61 bases by removing ten bases on the five prime end and three bases on the three prime end of each sequence. For 454-generated sequences, ten bases were removed on the five prime end only. The Illumina sequences were aligned onto the blueberry draft genome assembly [15] using TopHat2 [19] and Bowtie2 [20] using default parameters, except for the maximum intron size parameter, which was set to 6,000 bases consistent with typical intron size distributions for plant genes. 454 sequences were aligned onto the reference genome sequence using GMAP [21] with default parameters except for intron length, which was set to 6,000 bp.

### Data Availability

All sequence data are available in the Short Read Archive under accessions listed in the previous sections. Files contained alignments, RNA-Seq coverage graphs, and output from TopHat2 are available from a publicly accessible IGBQuickLoad site (http://www.igbquickload.org/blueberry) configured to serve data for visualization in Integrated Genome Browser. Data files including gene models and related annotations are available from a git source code repository at http://www.bitbucket.org/lorainelab/blueberrygenome. The git source code repository also contains R code used to analyze data.

### Gene model generation, filtering, and protein sequence assignment

Cufflinks was used to generate transcript models using paired-end and single-end Illumina sequences alignments with maximum intron set to 6,000 bases. Three ab initio gene-finding programs (Augustus [22], GlimmerHMM [23], and GeneMark [24]) were used to generate gene models from genomic sequence. Arabidopsis trained parameters were used for GlimmerHMM and default parameters were used for Augustus and GeneMark. Because many of the resulting ab initio and Cufflinks-based genes models covered the same genomic regions, a step-wise filtering protocol (Supplemental Figure 1) was applied to reduce redundancy in the final gene set. First, all genes generated by Cufflinks were selected for inclusion in the final gene set, since these were predicted from the RNA-Seq expression data and thus the level of evidence supporting them was high. Any ab initio gene finder-predicted gene that overlapped one of the Cufflinks-predicted genes was eliminated from the candidate gene list. Next, all remaining candidate genes that were predicted by Genemark were considered. If a candidate Genemark gene had homology to a known protein (by BLASTX) or overlapped with an expressed sequence alignment, it was added to the final gene set and, as before, overlapping gene models were removed from the remaining candidate gene set. The process was repeated for genes predicted by Glimmer, Augustus, and GMAP. The selection order of priority for ab initio gene prediction programs was based on visual inspection of predicted gene models and consistency with alignments of full-length blueberry sequences available from GenBank. Protein coding sequences were detected in the gene models using the TAU program [25]. Gene models were formatted into a BED-detail (BED14) file (V_corymbosum_scaffold_May_2013.bed) in which field 4 listed the transcript (gene model) name, field 13 listed the gene name, and field 14 listed the name of the program that generated the gene model. Note that many genes predicted by Cufflinks were associated with multiple transcripts, typically products of alternative splicing. Thus, in the case of Cufflinks-predicted genes, the gene field contained a name such as CUFF.11 while the gene model field contained a name such as CUFF.11.1, the first reported transcript generated from gene CUFF.11. Gene models in bed format were added to the bitbucket repository http://bitbucket.com/lorainelab/blueberrygenomeinsubdirectoryGeneModelAnalysis/results.

### Functional annotation of gene models using BLASTX against nr

Sequences of spliced transcripts were searched against the non-redundant protein database from NCBI using a database that was downloaded in June of 2013. Virtual cDNA sequences obtained from the gene model annotations were searched against the non-redundant (nr) database using blastx. The resulting matches were used for annotation if the e value was 10^−4^ or better, the alignment length was at least 30 amino acids, and the percent identity was 30% or higher. For each gene model with at least one high-quality match meeting these criteria, an annotation string was generated containing the Genbank identifier and fasta header for the best-matching sequence, information about the blastx-generated alignment, and the program that was used to generate the gene model. The annotation string was transferred onto the corresponding blueberry gene model by replacing field 14 in the BED-detail file described above. Field 14 data for gene models that did not have high-quality matches meeting the criteria described above were not modified. The modified gene model file was added to the bitbucket repository subdirectory titled BlastxAnalysis/results as V_corymbosum_scaffold_May_2013_wDescr.bed.gz.

### Functional annotation of gene models using Blast2GO

Blast2Go version 2.7.0 (Build 05122013) was obtained via Java Web Start from http://blast2go.com. and used to associate GO terms and EC numbers with individual gene models from blueberry. BLASTP and InterProScan search results were loaded into Blast2GO and the Blast2GO function **Annotation > Perform Annotation Step** menu was used to perform GO annotation. BLASTP results were obtained by searching predicted blueberry protein sequences against the nr protein database using evalue cutoff of 10^−3^ and reporting a maximum of five “hit” sequences per query. IPRScan version 4.8 was run using default parameter settings. Databases current as of July 2013 were searched. MySQL databases (‘b2g_sep13’) from Blast2Go.com were downloaded and installed on a local server to enable faster processing. Blast2GO results were saved in plain text format (Supplemental Data File 2) and in Blast2GO format and added to the bitbucket repository http://bitbucket.com/lorainelab/blueberrygenome in a subdirectory named Blast2GO.

### Functional annotation of gene models using BLASTX against PlantCyc enzymes

Fasta-format sequence files containing amino acid sequences for enzymes in Plant Metabolic Network species-specific databases (e.g., AraCyc, GrapeCyc, etc.) were downloaded in July of 2013 and combined into one database. The blueberry cDNAs were used as queries in a BLASTX search of the combined database. Hits to PlantCyc enzymes were then filtered so that hits with at least 60% subject coverage, 45% identity or higher, and e-value 0.001 or less were retained. Details of how this was done are explained in Markdown file MakingAnnotationFiles.Rmd in the bitbucket repository for the paper. The best hit for each blueberry gene model was identified by e value and used to generate annotation text, which was inserted into field 14 of the gene models bed file and saved to subdirectory PlantCyc/results as V_corymosum_scaffold_May2013_wDescrPwy.bed. To enable keyword searching in Integrated Genome Browser, this annotation text included the best matching PlantCyc enzyme and a list of PlantCyc pathway identifiers when available.

## Analyses

### Building and filtering blueberry gene models

To characterize gene expression in blueberries, Illumina and 454/Roche sequencing of green and ripe blueberry cDNA was done, including a developmental time course experiment in which three biological replicates of five berry fruit stages were sequenced using 100 base, single-end Illumina HiSeq sequencing. Table I summarizes the sequencing strategies used and amount of sequence obtained, which totaled around 800 million sequence RNA-Seq “reads” and more than 75 billion bases. Most sequence data were from the developmental time course experiment, which surveyed berry samples collected from three individuals of the O’Neal cultivar, an early ripening variety of southern highbush blueberry that is widely grown in North Carolina and other southern US states. The second largest RNA-Seq data set was from paired-end, 75 base pair Illumina sequencing of ripe and green berries collected from the O’Neal, Arlen, Lenoir and Pamlico southern highbush cultivars. Analysis of genetic differences between the cultivars will be described elsewhere, but the data were included here to enable a more complete transcriptome assembly and analysis. In addition, two full plates of 454 sequencing of ripe and unripe berries from the O’Neal cultivar were done using berries collected during spring and summer of 2009 and 2010. Because of the abundance of data and availability of easy-to-use software for generating gene models from Illumina-based RNA-Seq data, we used the Illumina sequence data to generate a genome-guided transcriptome assembly and reserved the 454 data for gene model validation and assessment.

**Table I.** Summary of sequencing strategies and sequences obtained. The number of sequences obtained from each sequencing strategy is shown.

| Sequencing Strategy | Samples | No. of reads (millions) | Bases (billions) |
| --- | --- | --- | --- |
| Single-end Illumina, 100 bp reads | Five fruit stages from O'Neal cultivar | 485 | 48.6 |
| Paired-end Illumina, 76 bp reads | Ripe, unripe berries from O'Neal, Lenoir, Ozark blue, and Pamlico cultivars | 377 | 28.3 |
| 454 | Ripe and unripe berries from 2009, 2010 harvests, O'Neal cultivar | 2.16 | 0.73 |
| Total |  | 864.16 | 77.63 |

To generate berry gene models from the RNA-Seq data, the Illumina sequence reads were aligned onto the May 2013 reference blueberry genome using the spliced alignment program TopHat [19] and then merged into gene models using Cufflinks [26]. This step produced 64,666 transcript models representing 57,925 genes. Of the multi-exon genes, 24% were predicted to generate multiple transcript variants due to alternative promoter use or alternative splicing. To assess how realistic this level of transcript variation was, we compared the frequency of alternative transcripts in blueberry to alternative transcription rates in Arabidopsis and soybean, using the genome annotations and assembly releases that were available in July 2013 for those species. The alternative transcription rates among multi-exon genes for soybean and Arabidopsis were 25% and 26% respectively; these rates were similar to blueberry, indicating that the frequency of transcript variation found in the blueberry gene models was reasonable and not likely to be an artifact of incorrect transcript model assembly.

Because the Cufflinks gene models were based on berry fruit RNA-Seq data, some genes that were expressed primarily in other sample types (e.g., roots) might be missed. To complement the Cufflinks-generated gene models, ab initio gene prediction programs were used to generate additional gene models from the reference genomic sequence (Supplemental Figure 1A). The ab initio gene finding programs generated more than 185,000 gene models, of which 75% overlapped with genes identified in the RNA-Seq data by Cufflinks and therefore were likely to be redundant. To eliminate duplicates and create a non-redundant collection of blueberry gene models, a stepwise filtering protocol was applied (Supplemental Figure 1B) that retained gene models based on their relative level of evidence. To start with, all gene models based on the aligned RNA-Seq data were added to the non-redundant set. Next, gene models with homology to known proteins were selected. Following this step, gene models were kept according to which ab initio gene finding program produced them. We found through manual inspection in Integrated Genome Browser (IGB) that some programs produced more realistic models when visually compared to the 454 alignments, and so we configured the filtering pipeline based on these observations. Note that this final filtering step could be further optimized through systematic comparative evaluation of ab initio models to RNA-Seq based gene models, but due to the limitations of time and data availability, we did not do this. Ultimately, the final non-redundant set of gene models included 70,581 gene models and 63,840 genes, of which most were based on the RNA-Seq data (Table II).

**Figure 1.**
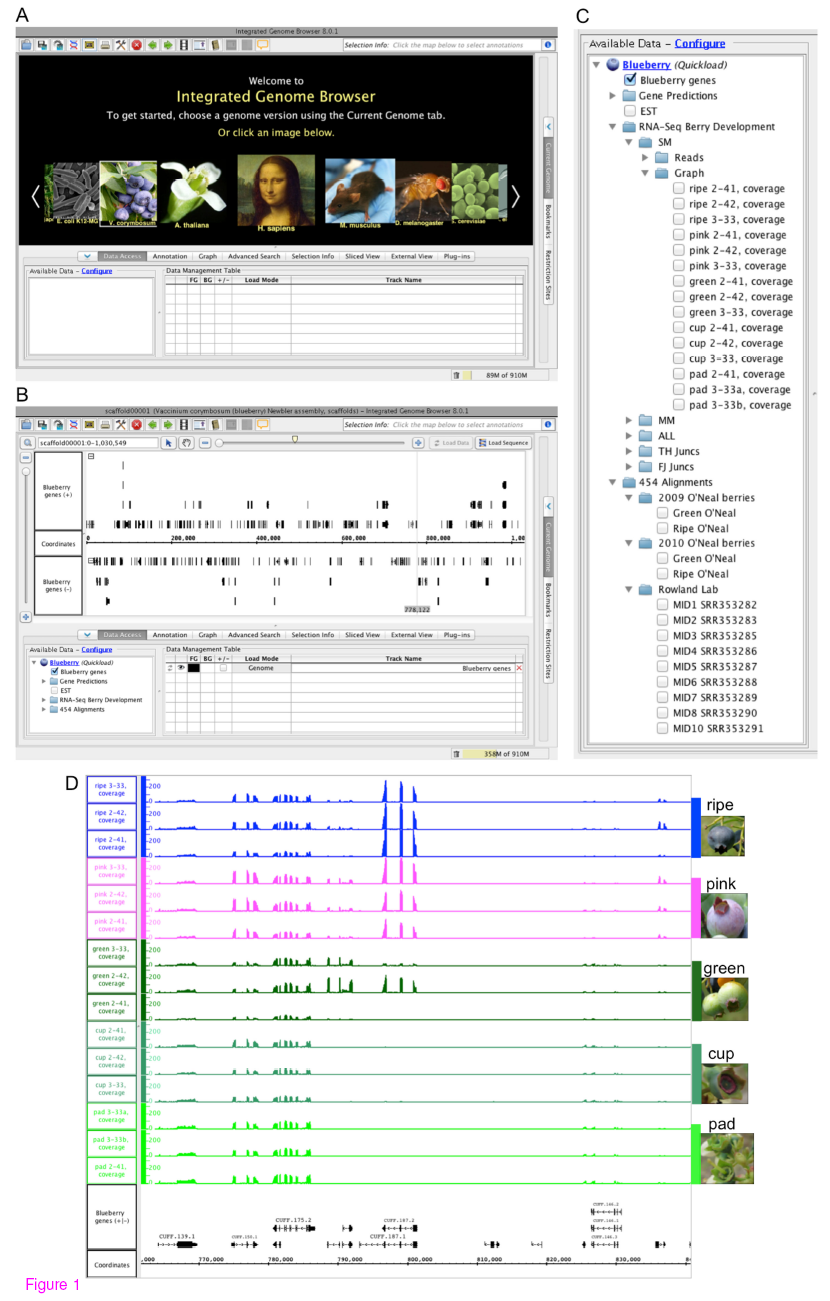
Blueberry genes in Integrated Genome Browser. **(A)** IGB start screen showing shortcut to the blueberry genome. (**B**) Clicking the image labeled *Vaccinium corymbosum* triggers loading of blueberry genome assembly May 2013 and the non-redundant gene set. (**C**) Blueberry data sets available in IGB. (**D**) Coverage graphs showing read density on a region of scaffold00001.

**Table II.** Genes predicted by Cufflinks RNA-Seq analysis or ab initio gene-finding programs. The number of genes in the non-redundant gene set is shown.

| Method | Genes |
| --- | --- |
| Cufflinks (RNA-Seq) | 56,087 |
| GeneMark.hmm | 4,794 |
| Augustus | 1,745 |
| GlimmerHMM | 933 |
| 454Scaffolds (454 RNA-Seq) | 281 |

Next, an open read frame (ORF)-finding program (TAU, [25]) was used to identify and annotate a conceptual translation for each gene model. Altogether, there were 57,079 gene models from 51,515 genes that could be annotated with a protein sequence of at least 30 residues, the expected lower range of protein sizes based on Arabidopsis annotations. The remaining transcript models for which no longer protein sequence could be predicted had a broad range of sizes and exon numbers, ranging from short, single-exon genes to longer models with multiple exons, suggesting that in many cases, the gene models were simply incomplete or the sequence itself contained insertions, deletions or other problems that stymied conceptual translation. Another possibility was that many of these genes were non-coding genes whose primary products were RNA and not protein. A third possibility was that many of these were actually pseudogenes. An analysis of Arabidopsis RNA-Seq data [27] found that many non-coding genes, including both pseudogenes and non-protein coding genes, were detectable in libraries that were prepared using the same protocol as with the blueberry RNA-Seq libraries, and so these latter two possibilities may have been correct in many cases.

It is important for researchers to have access to the blueberry data in formats that facilitate exploration, visualization, and analysis of individual sequences, genes, and gene families. To enable better use of the data, the blueberry genome assembly, annotations, and RNA-Seq data have been made available in the Integrated Genome Browser, an open source, desktop genome visualization platform that is freely available for download from http://www.bioviz.org. IGB uses a companion IGBQuickLoad Web site to distribute data sets in formats that enable both interactive visualization and larger-scale analysis. For example, users can download data sets in formats that are amenable to bioinformatic analysis, such as the BAM and bed formats, or instead can simply view a listing of the data sets in IGB and open them there. To open the blueberry genome in IGB, users first select the blueberry image shortcut from the IGB start screen (Figure 1). Clicking the blueberry image triggers loading of the non-redundant gene data set described above along with a listing of blueberry genome scaffold and their sizes (in the **Current Genome** tab) and a catalog of available data sets (in the **Data Access** tab). As of this writing, the available data sets included the Illumina and 454 RNA-Seq data described here as well as reference data sets from other studies, including a 454-based sequencing data set that included multiple sample types and berry fruit ripening stages [16]. Thus IGB allows researchers to view and explore multiple data sets from diverse sources, not just the current study.

Once the genes are loaded, users can search for genes by name or by annotation key word using the **Quick Search** (top left) or **Advanced Search** tab, which also enables searching for sequence motifs. Gene models are associated with descriptive text assembled from homology searches against blast databases from NCBI and PlantCyc (described below), but users can investigate individual gene models in greater detail using the blast features available through IGB. Right-clicking a gene model displays a menu that displays options to run blastx or blastp searches against the nr protein database at NCBI. Selecting one of these options opens a Web browser window, which shows results from the search. In addition to the blast search feature, the right-click menu (also called a “context menu”) offers the option to show the sequence of a genome model in a separate window, which in turn enables selecting and copying the protein sequence. Using this feature, users can easily copy and paste the protein sequence into other Web-based search and analysis tools, such as the InterproScan search tool, which can identify conserved motifs in protein sequences.

Users can also load the RNA-Seq data sets and use the observed pattern of expression to gain insights into the function of individual genes. To illustrate, the image shown in Figure 1 depicts a region from scaffold00001 showing gene models and RNA-Seq coverage graphs from five berry developmental stages. Because the overall read counts for each track were similar, one can visually estimate and compare expression levels for genes provided the coverage graphs are put onto the same scale, which is possible using the IGB **Graph** tab. In this example, comparing graph heights between stages highlights ripening related expression of gene CUFF.187, which is homologous to genes in many other plant species but has no known function. The stage-specific expression of CUFF.187 demonstrates the power and usefulness of visualizing gene models alongside expression data within a visualization environment that also enables rapid exploration of genes and their functions.

### Functional annotation of blueberry gene models and comparison with other plant species

To identify putative functions of newly annotated blueberry genes, we searched for homologous proteins using blastx searches of the non-redundant nr database from GenBank. We found that most blueberry genes (58%) had significant homology (e value ≤ 0.001 and percent identify ≥ 30%) with at least one protein in the nr protein database. Typically, the highest scoring, best match to a blueberry protein was a protein from the wine grape *Vitis vinifera* (Figure 2A). The abundance of best-scoring hits from grape was probably due to two factors: the similarity between grape and blueberry [28] and the large number of grape sequences that were present in the nr database at the time this analysis was done. The nr database contained nearly 80,000 proteins from grape, many of which were conceptual translations from the sequenced grape genome, which has been available since 2007 [29]. The tea plant *Camellia sinensis*, also represented in Figure 2A, is more closely related to blueberry than grape, but had fewer best hits overall primarily because there were fewer tea sequences in the databases. We found that in general, if a blueberry gene had a match to a tea protein, the tea protein match had a better score and higher percent identity than the corresponding best match from grape.

**Figure 2.**
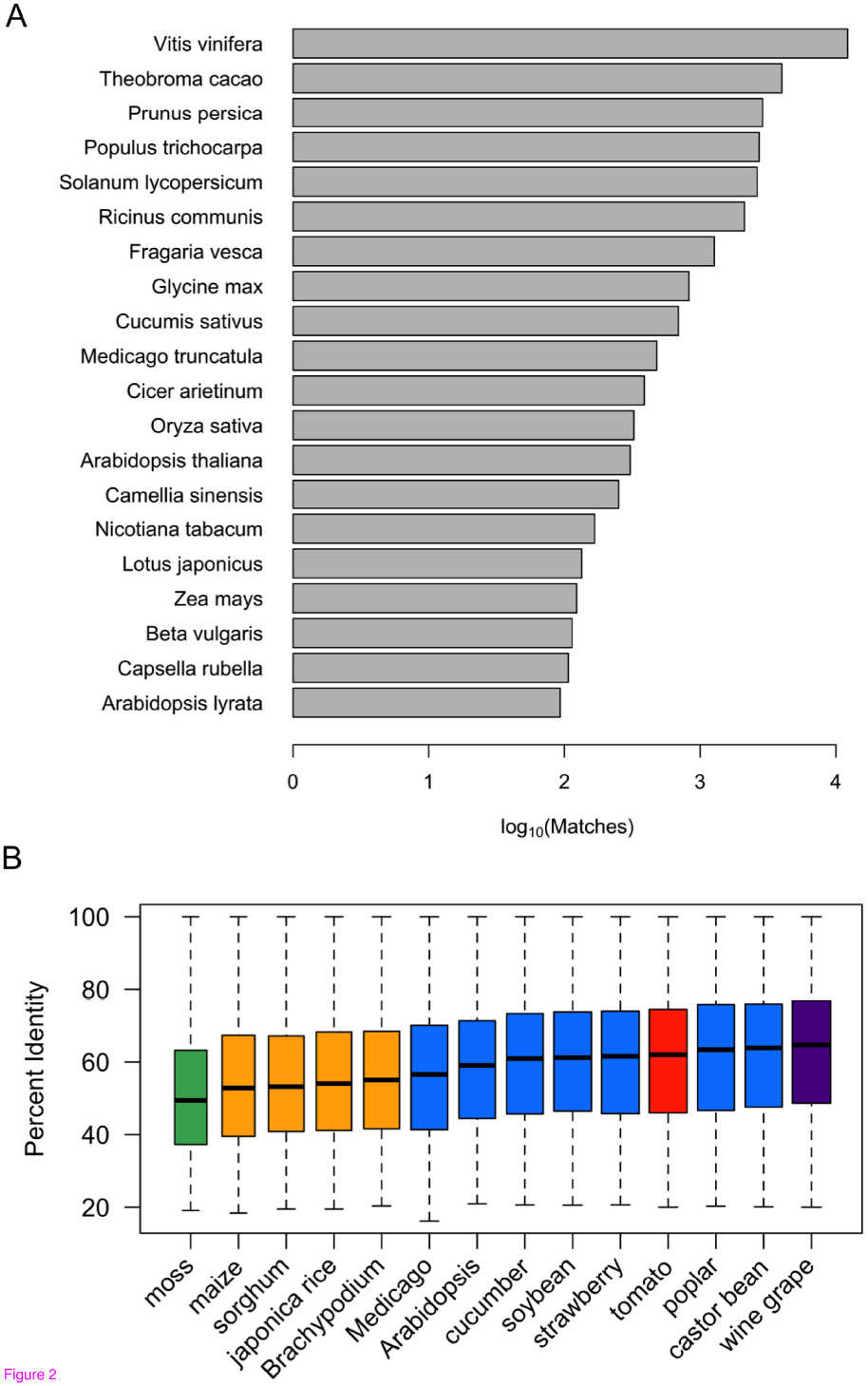
Results from searching nr and RefSeq protein databases. (A) Number of best-matching proteins by species. Blueberry transcript sequences were searched against the non-redundant protein database from NCBI and the best-matching protein for each blueberry sequence was identified. The plot shows the number of blueberry genes whose best matching protein was from the indicated species. (B) **Distribution of percent identity scores by plant RefSeq database.** Blueberry transcript sequences were used to search RefSeq protein databases for plants with close-to-complete, annotated genomes. Boxplots show the distribution of percent identity scores by species. RefSeq databases include grape (*Vitis vinifera*), castor bean (*Ricinis communus*), poplar (*Populus trichocarpa*), tomato (*Solanum lycopersicum*, Sl), strawberry (*Fragaria vesca*), soybean (*Glycine max*), cucumber (*Cucumus sativus*), Arabidopsis (*Arabidopsis thaliana*), Medicago (*Medicago trunculata*), Brachypodium (*Brachypodium distachyon*), rice (*Oryza sativa*), sorghum (*Sorghum bicolor*), corn (*Zea mays*), and a moss (*Selaginella moellendorffii*).

Nonetheless, these results prompted us to explore relationships between blueberry and other plant genomes. For this, we used plant-specific RefSeq databases from GenBank, including databases for grape, tomato, Arabidopsis, and several other plant species where a close-to-complete, well-annotated proteome was available [30]. To characterize relationships between blueberry and these other plants, we searched the blueberry transcripts against these close-to-complete plant RefSeq databases and identified the best scoring protein from each genome for each blueberry gene model. Figure 2B shows the distribution of percent identity scores obtained for the best-matching sequences from each species. Grape proteins had the highest median percent identity scores, followed by poplar, castor bean, and tomato. Blueberry and other *Ericales* species are part of the asterid clade of flowering plants, and are more closely related phylogenetically to tomato than to rosids *Populus trichocarpa* (poplar) and *Ricinus communis* (castor oil plant) [31]. The blast results do not contradict this relationship but instead highlight how sequence similarity may reflect similarities in physical or biochemical characteristics. Grape and blueberry are deciduous, berry-producing plants with long generation times. Poplar is also a deciduous, woody plant with a long generation time. Biochemical or morphological similarities that could explain the similarity between blueberry and castor bean are less obvious, however. These results suggest that in-depth comparison between grape, blueberry, and castor bean seed transcriptomes could lead to new insights into developmental programs at work in berries of the three species.

### Blast2GO annotation

Blast2GO is a popular annotation platform that uses results from homology searches to associate sequence with Gene Ontology (GO) terms and other functional annotations [32], a critical step toward enabling analysis of high-throughput gene expression studies. To facilitate analysis of the blueberry developmental time course data and also provide a resource for other blueberry researchers, we used Blast2GO to annotate the blueberry proteins with GO terms. Blueberry sequences were searched against the nr protein database and the InterPro database [33] and the results were imported into the Blast2GO program’s graphical user interface, which assigned GO terms to 18,143 transcripts representing 15,308 genes. Plots summarizing GO terms and the number of gene products assigned to each term are shown in Figure 3, and a file containing the GO term mappings is provided as Supplemental Data File S3. Similar to Arabidopsis [34] and other plants, a large number of genes were annotated with terms related to transcription factors. Around 1,300 transcripts received GO annotation “DNA binding,” of which around 400 were also were annotated with the more specific GO term “sequence-specific DNA binding transcription factor activity.” These sequences likely represented expressed transcription factors involved in regulation of gene expression during fruit development and ripening. More than 500 proteins were associated with terms related to primary and secondary metabolism, lipid metabolic processes, or carbohydrate metabolic processes, reflecting the dynamic processes underway in fruit development. Thus the GO annotation identified a large set of candidate genes likely to be involved in regulation and synthesis of bioactive secondary metabolites.

**Figure 3.**
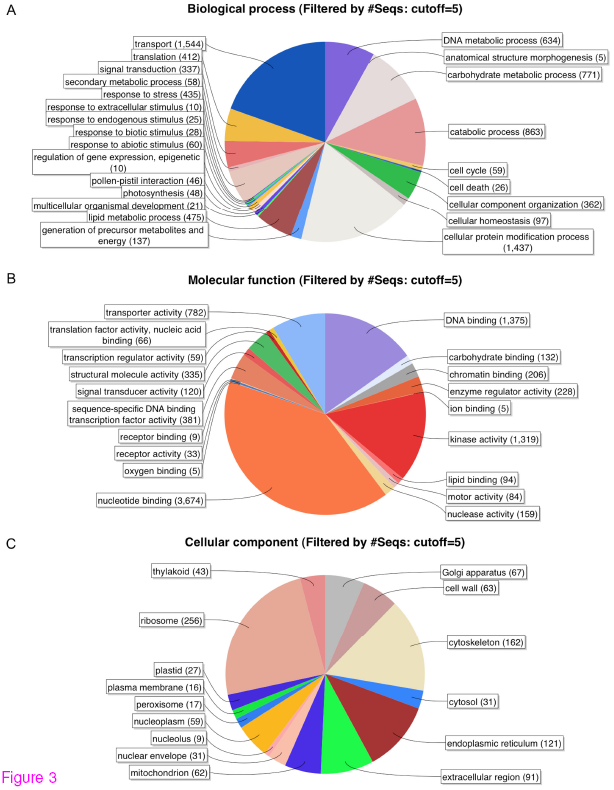
Gene Ontology annotations for the three sub-trees of GO. (A) Biological process Molecular Function (C) Cellular Component. Numbers indicate the number of protein-coding blueberry transcripts assigned to each category. Plots were made using Blast2GO. The number of genes assigned to each category can be obtained from Supplemental Data File S3.

### Metabolic pathway annotation using PlantCyc enzyme database

The PlantCyc database is a collection of curated and computationally-predicted enzymes, enzymatic reactions, and metabolic pathways for 18 plant species, including grape, Arabidopsis, cassava, poplar, and rice [35]. The PlantCyc databases are closely tied with the Pathway Tools software package [36], a visualization and database system for pathways and biochemical reactions. At the time of this writing, the pathways database for *Vitis vinifera* (GrapeCyc version 3.0) was one of the most complete, containing annotations for 432 pathways. The similarity between grape and blueberry sequences suggested that comparing the blueberry sequences to enzymes in GrapeCyc and other PlantCyc databases would identify candidate genes encoding enzymes of primary or secondary metabolism. To identify enzymatic functions for blueberry genes, the BLASTX algorithm was used to search for matching PlantCyc sequences. To eliminate matches arising from alignments between domains that occur in many different sequences (e.g., ATP-binding cassette), only matches that covered at least 60% of the subject sequence with at least 45% identity were considered. Under these criteria, transcripts from more than 7,100 blueberry genes were found to match at least one PlantCyc enzyme sequence, and there were over 450 pathways from PlantCyc that had at least one enzyme matching a blueberry sequence. As before, grape proteins were typically the best matches for blueberry sequences.

### In-depth analysis of ethylene biosynthetic pathway gene expression

Ethylene is a gaseous plant hormone that controls many aspects of plant development and physiology, especially ripening. In climacteric fruits such as tomato and banana, a burst of ethylene biosynthesis triggers ripening, and post-harvest treatment with exogenous ethylene can control ripening onset and progression. By contrast, blueberries do not appear to undergo a burst of ethylene synthesis prior to ripening, and post-harvest application of ethylene has far less effect on the ripening process. Nonetheless, ethylene can influence aspects of flowering and ripening, as shown by experiments with ethephon, a horticultural chemical that when applied to foliage is enzymatically converted to ethylene. When applied to blueberry plants during the harvest, ethephon accelerates and synchronizes ripening, and when applied in the fall, ethephon delays flowering the following spring and increases the number of flower buds (reviewed in [37]). This suggests that ethylene is important in flowering and fruit development but its role is likely to be very different than in climacteric fruits.

Ethylene is synthesized from L-methionine via three reactions that are catalyzed by SAM synthase (methionine adenosyltransferase), ACC synthase (1-aminocyclopropane-1-carboxylate synthase), and ACC oxidase (1-aminocyclopropane-1-carboxylate oxidase). In most plants, large multi-gene families encode all three enzymes. The PlantCyc database reports that the grape genome contains 34 SAM synthase genes, 20 ACC synthase genes, and 12 ACC oxidase genes. Consistent with this, the PlantCyc-based annotation of blueberry identified similarly large numbers of genes encoding ethylene biosynthetic genes, including 22 genes encoding SAM synthase, six genes for ACC synthase, and seven genes encoding ACC oxidase (Supplemental Figure 2). Interestingly, the expression patterns for SAM synthases were highly variable and at least one SAM synthase gene was highly expressed (more than 1,000 RPKM) in each stage of berry fruit development and ripening. Expression of genes encoding ACC synthase, a key control point for ethylene and ripening in tomato, was also highly variable. Some genes encoding ACC synthases were expressed at very low levels (close to 0 RPKM) while others were expressed at 200 RPKM or higher, which was around the 95^th^ percentile of gene expression as measured in RPKM. Two ACC oxidase genes were expressed at 150 RPKM or higher at each stage, while others were expressed at much lower levels. Interestingly, one ACC oxidase gene (CUFF.81159) was highly expressed during ripening and reached more than 3,000 RPKM in ripe fruit. The uniformly high expression (>150 RKPM) of ethylene biosynthesis genes at every stage, combined with the extremely high expression of ACC oxidase gene CUFF.81159, suggested that ethylene is produced throughout berry fruit development and that these levels are likely to peak in ripe fruit.

### Anthocyanin biosynthetic pathways

Anthocyanins are the 3-*O*-glycosylated forms of anthocyanidins, which consist of a polyphenolic ring substituted with –H, -OH, and –OCH3 groups at different positions. The substituted group and its location on the polyphenolic ring determine the type of anthocyanidin and may also dictate aspects of biological activity. In blueberry, the anthocyanins containing malvidin, delphinidin, and peonidin aglycones are especially abundant by weight [4]. Another level of diversity arises from the type of sugars attached, and these sugar groups may influence bioavailability in the mammalian digestive tract [38, 39]. Thus it is of interest to identify enzymes that catalyze steps in anthocyanin biosynthesis and profile their expression pattern and relative abundance in ripe fruit.

Comparison with PlantCyc enzymes identified 31 genes with homology to enzymes involved in anthocyanin synthesis, including two genes (CUFF.20951 and CUFF.43605) that were highly expressed in pink and ripe fruit in comparison to other genes in the pathway (Figure 4). CUFF.20951 was the most highly expressed; it reached more than 12,000 RPKM in pink and ripe fruit, up from around 2,000 RPKM in green fruit. The best matching PlantCyc enzyme for CUFF.20951 was an enzyme from poplar that was annotated as anthocyanidin-3-*O*-glucosyltransferase, which transfers a glucose sugar onto the 3-*O*-position of the anthocyandin polyphenolic ring. Searching the nr database using blastp identified even better matches, including predicted flavonoid galactosyl and glucosyltransferases from a variety of plant species. Based on its homology to other sequences, the CUFF.20951 protein likely is involved in glycosylation of flavonoids, but its preferred aglycone and sugar substrates are impossible to determined based on sequence data alone. However, judging from the extremely high expression of this gene in blueberry, the CUFF.20951 protein is likely to be responsible for a high percentage of anthocyanin (or other flavonoid) production in ripe berry fruit.

**Figure 4.**
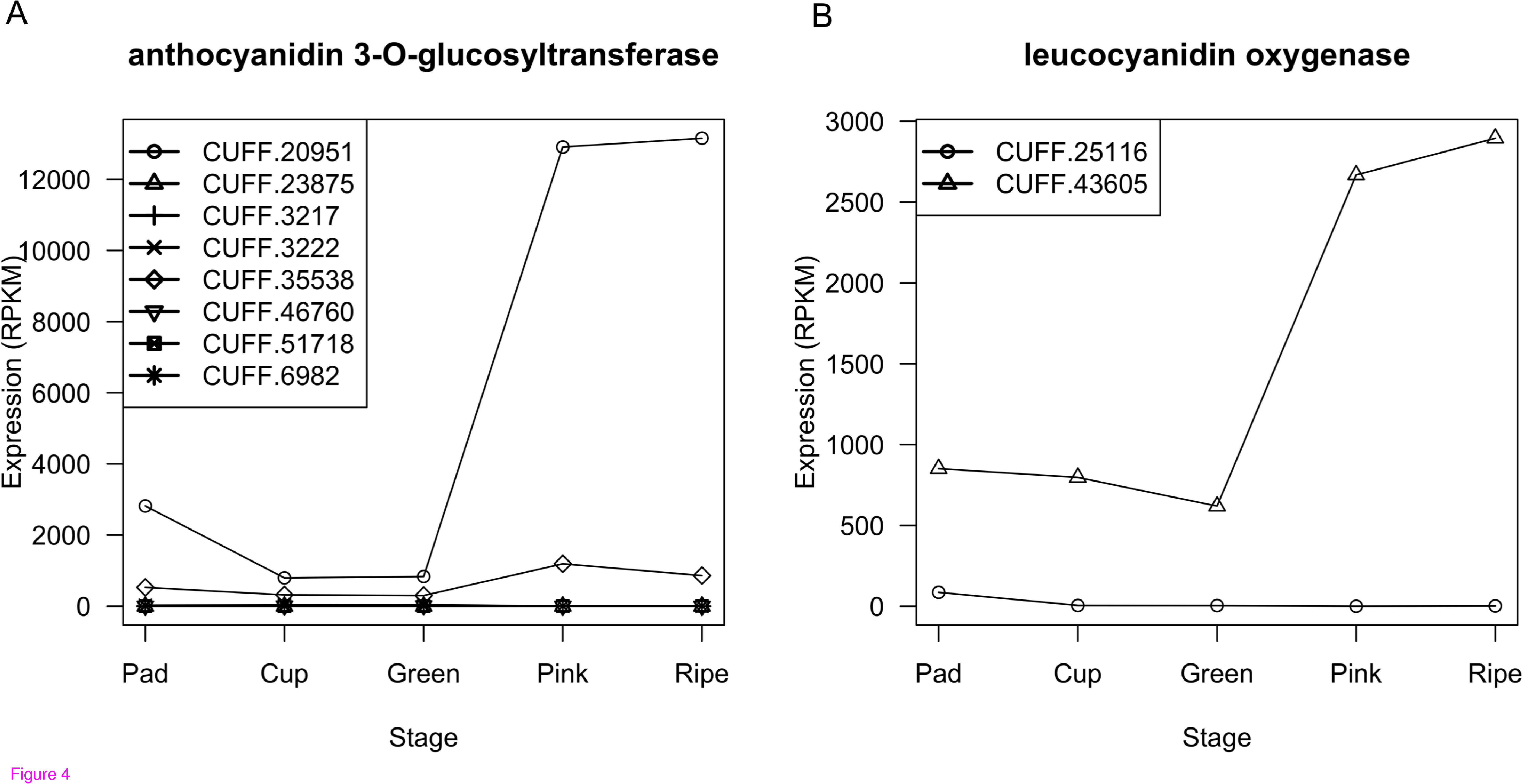
Expression pattern of anthocyanin biosynthetic genes. Expression pattern for genes annotated to enzymatic reactions in anthocyanin-related pathways in PlantCyc are shown. (**A**) Genes encoding leucoanthocyanidin dioxygenase, catalyzing EC 1.14.11.19, dihydroquercetin synthesis from leucoanthocyanidin or cyaniding synthesis from leucoanthocyanidin (**B**) Genes for enzymes catalyzing 3-O-glucosylation of delphinidin (EC 2.4.1.115). Both are annotated to PlantCyc pathway anthocyaninin biosynthesis (delphinidin 3-O-glucoside).

The other highly expressed gene was CUFF.43605, which reached a peak of around 2,000 RPKM in pink and ripe fruit, up from around 900 RPKM in the earlier stages. The best PlantCyc match for CUFF.43605 was a protein from *Brassica rapa* (Bra019350, 78% identity) annotated as a leucocyanidin oxygenase, also called anthocyanidin synthase (ANS) or leucocyanidin dioxygenase (LDOX). Homologous enzymes from Arabidopsis [40, 41] and rice [42] convert leucoanthocyanidin to anthocyanidins, precursors for anthocyanins, but they can also catalyze formation of other bioactive flavonoid precursors, notably dihydroquercetin, the precursor of quercetin. Anthocyanins and quercetin are both abundant in berries, but quercetin has greater bioavailability and therefore may be a more potent bioactive in berry fruit [43, 44]. If the preferred end product of the CUFF.43605 reaction is indeed dihydroquercetin, then its remarkably high expression likely has a positive effect on quercetin concentration in berry fruit. However, if its major end product is cyanidin, then it likely acts to decrease quercetin levels by consuming leucoanthocyanidin, the dihydroquercetin precursor. As with CUFF.20951, sequence and expression analysis alone is likely insufficient to distinguish these possibilities. Nonetheless, the high expression of CUFF.43605 makes this gene a fruitful candidate for investigating genetic control of anthocyanin and quercetin abundance in berries.

### Other pathways – bixin, and dhurrin

Bixin is the primary component of annatto, a commonly used food dye collected from the seeds of *Bixa orellano*, which grows in the tropics and is also known as the lipstick plant. Annatto is used in folk medicine and for body decoration and has also been investigated as a plant-based treatment for diabetes [45], cancer [46], and microbial infections [47]. We found that four blueberry genes matched two of three annotated enzymes of the pathway (Supplemental Figure 3). Expression of all four genes peaked in the mature green stage of fruit development, around the time when seeds were developing. Grape seed contains bixin [48], and, as discussed previously, the grape and blueberry proteomes were remarkably similar. These results suggested that blueberries may contain a bixin-like compound. .

Blueberry genes similar to enzymes from a potentially harmful pathway were also found. Blast analysis identified putative blueberry homologs for each of three biosynthetic enzymes involved in synthesis of dhurrin in sorghum [49-51]; the blueberry genes shared between 45 and 60% identity with their putative homologs from sorghum. Dhurrin, which reaches up to 10% dry weight in sorghum seedlings, is one of a large class of cyanogenic glycoside (CG) defense compounds plants synthesize from amino acid precursors as a form of chemical warfare against insects and other herbivores [52, 53]. Stored as inactive glycosides, mechanical damage to cells (such as from chewing) activates endogenous glycosidases that remove the sugar group, triggering production of toxic hydrogen cyanide (HCN) due to instability of the sugar-free aglycone or to the activity of other catabolic enzymes. Interestingly, the putative blueberry homologs were most highly expressed in the green fruit stages (Supplemental Figure 3D), suggesting that green berries synthesize a cyanogenic glycoside that discourages insects and mammals from eating unripe berries.

Homology searches also identified putative berry homologs of enzymes involved in two CG catabolic pathways, one that removes the glycosyl group from the CG leading to production of HCN, and another pathway that detoxifies CGs by converting them to harmless byproducts. Seven genes had significant similarity to enzymes in the cyanogenic catabolic pathway, including four genes resembling dhurrinase, which deglycosylates dhurrin, and three genes that were similar to hydroxynitrile lyase, which catabolizes the dhurrin aglycone to HCN. Similar to the biosynthetic enzymes, these catabolic, cyanogenic enzymes were most highly expressed in unripe fruit. We also identified a blueberry candidate gene encoding nitrilase 4, an enzyme that detoxifies dhurrin and possibly other cyanogenic glycosides by converting them to aspartic acid and asparagine [54]. Searching with nitrilase 4 from *Arabidopsis thaliana* (AtNIT4, AT5G22300) identified blueberry gene CUFF.32314, which shared 83% identity with the Arabidopsis protein. Plant nitrilase 4 enzymes are highly conserved [54], typically sharing 60 to 70% identify at the amino acid level, suggesting that CUFF.32314 indeed encodes nitrilase 4. The putative blueberry nitrilase CUFF.32314 gene was highly expressed in young fruit and ripe fruit, reaching more than 70 RPKM in ripe berries, but had much lower expression (<40 RPKM) in mature green berries. Thus the expression profile of the candidate gene involved in CG detoxification was roughly complementary to profiles of candidate genes involved in CG synthesis and cyanide release. This suggests immature green berries produce CG compounds that discourage herbivory, but then berry NAT4-like genes de-toxify these CGs as part of the ripening process.

### Differential gene expression during blueberry development and ripening

Fruit development and ripening are dynamic processes involving cell division, expansion, differentiation, synthesis of pigments and other secondary metabolites, as well as import of sugars into the fruit (reviewed in [55]). To better understand these processes in blueberry, we used the developmental time course RNA-Seq data to examine global patterns of gene expression throughout fruit development, maturation, and ripening. Statistical analysis of gene expression identified nearly 19,000 genes that were differentially expressed between at least two stages, even when a relatively stringent false discovery rate cutoff of 0.001, corresponding to roughly 1 in 1,000 false discoveries, was applied. Table III reports the number of genes that were differentially expressed between stages, broken down by up- and down-regulation. Except for the pink versus ripe fruit comparison, there were more up-regulated than down-regulated genes in every comparison between a later stage and an earlier stage, reflecting the greater diversity of genes that were expressed in the later stages as tissues continued to develop and differentiate and as the seeds formed within the fruit. Viewed globally, the gene expression changes that occurred during progression of fruit growth, maturation, and ripening were extensive and dynamic.

**Table III.** Up- and down-regulated genes between sample types. Each cell in the table represents a comparison between two sample types. Cells above the diagonal report the number of genes that were up regulated in the later of the two compared stages. Cells below an empty cell report the number of genes that were down regulated in the later of the two compared stages. Cutoff for differential expression was false discovery rate of 0.001. Stages are reported in order of earlier (Pad) to later (Ripe).

|  | Pad | Cup | MG | Pink | Ripe |
| --- | --- | --- | --- | --- | --- |
| Pad | - | 566 | 2,390 | 6,010 | 6,077 |
| Cup | 526 | - | 1,221 | 5,270 | 5,003 |
| MG | 1,981 | 919 | - | 3,583 | 3,726 |
| Pink | 6,765 | 6,594 | 3,038 | - | 18 |
| Ripe | 6,821 | 6,466 | 5,159 | 56 | - |

Figure 5 reports the number of genes that were differentially expressed (DE) between adjacent stages and highlight the physiological transitions as fruits enlarge from small, pad-like structures on the base of the pedicel to round, mature berries harboring seeds, flesh, and sugars. The most similar of the adjacent stages were the pink and ripe fruit, collected 14 days apart. Fewer than 100 genes were DE between pink and ripe fruit. Of these, most (75%) were down regulated. The up regulated genes included genes of unknown function and genes involved in enzymatic activity, such as UDP-glucosyltransferases, an alcohol dehydrogenase, and hormone biosynthetic genes. Down-regulated genes included many enzymes, such as the putative dhurrin biosynthetic enzymes, as well as other genes of unknown function. By contrast, the pad and cup stages, which were collected 10 days apart, had more than ten times as many DE genes. Gene Ontology enrichment analysis of genes differentially expressed between the pad and cup stages found that a significant number of DE genes were annotated with terms related to transcriptional regulation, regulation of enzymatic activity, and cell wall biosynthesis (Supplemental Data File S4). The type and high number of DE genes in the pad-to-cup versus the pink-to-ripe comparison reflects the extent of developmental changes underway during early, rapid growth stages of berry development.

**Figure 5.**
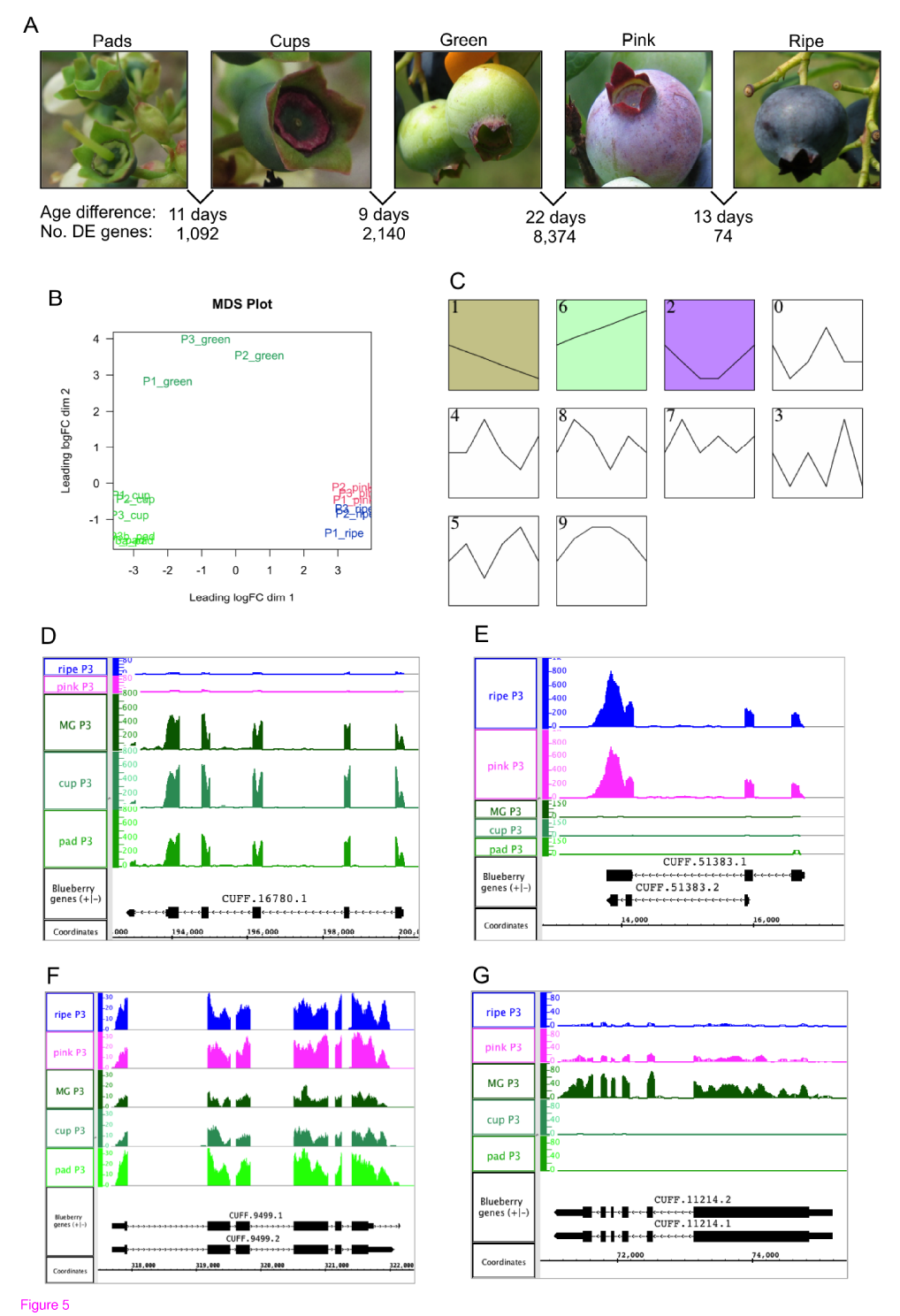
RNA-Seq analysis of gene expression changes during berry fruit development and ripening. **(A)** Photographs of berries exemplifying five stages of berry fruit development and ripening and the number of differentially expressed genes between adjacent stages. (**B**) Multi-dimensional scaling plot of expression values from five sample types and three plants. Axes represent first and second principal components and therefore have no units. Samples that are near each other in the two-dimensional space are more similar with respect to gene expression. Gene expression profiles detected by clustering program Short Time-series Expression Minor (STEM). Clusters 0 through 9 are listed in order of statistical significance, with clusters 1, 6, 2, and 0 being the most significant. **(D-G)** Cufflinks-predicted genes from clusters 1, 6, 2, and 0 encoding proteins similar to (**D**) leucoanthocyanidin reductase (**E**) MYB-like transcription factor up-regulated in ripening fruit. (**F**) tryptophan synthase (indole-salvaging) (**G**) abscisic acid signaling transcription factor ABI3. In (**D** – **G**), graphs represent the number of reads overlapping genomic positions indicated on the coordinates axis. RNA-Seq coverage graphs from P3 (plant 3-33) are shown.

However, the transition from fully rounded green fruit to pink fruit involved by far the largest number of differences. The chief physical difference between the two stages was color; pink berries had some reddish color indicating the onset of ripening, but other than this, green and pink berries were similar. Gene Ontology enrichment analysis identified around 40 terms as significantly enriched among genes differentially expressed between the green and pink fruit. Two terms were related to photosynthesis and likely reflected down-regulation of photosynthetic functions, despite the fact that when the pink berries were collected, they typically were only partly pigmented and much of the berry surface was still green. Of 43 genes annotated to the term “photosynthesis,” 27 were differentially expressed and all were down regulated in pink fruit. Other significant terms were related to metabolic pathway functions, including the terms metabolic process, catalytic activity, catabolic process, hydrolase activity, carbohydrate metabolic process, lipid metabolic process, transferase activity, and biosynthetic process. There was also enrichment in functions related to transcriptional regulation of gene expression, including chromatin binding, nucleic acid binding, sequence-specific DNA binding transcription factor activity, and signal transduction.

Multi-dimensional scaling, similar to principle components analysis, was used to cluster samples according to their overall similarity of gene expression pattern (Figure 5B). Samples from similar stages formed clusters. Pink and ripe fruit formed a cluster, cup and pad stages formed another cluster, and mature green fruit formed another cluster separate from the others. Interestingly, one ripe fruit sample (plant 2-41, P1) clustered slightly distant from the other ripe fruit samples. Visualization of RNA-Seq coverage graphs in IGB showed that some genes were up or down regulated in this sample relative to the other ripe fruit samples. Genes that were down regulated in P1 relative to the other ripe berry samples had functions related to cell wall degradation or modification, whereas genes that were up regulated had functions related to stress responses. This suggests that the P1 ripe berry sample was undergoing a stress relative to the other samples. Nonetheless, the possible presence of a possible confounding factor related to stress did not significantly reduce power to identify differentially expressed genes between samples types.

A time series clustering program (STEM [56]) was used to identify groups of genes that varied in concert over time. Scaled, averaged expression values for genes found to be DE between any two fruit stages were provided as inputs to the STEM program, which identified several gene expression profiles depicted graphically in Figure 5C. Two of the most statistically significant profiles had complementary patterns (clusters labeled #1 and #6). The “early high” profile (#1 in Figure 5C) contained genes with high expression in early fruit stages and lower expression in later stages. Gene Ontology enrichment analysis showed that this early high cluster contained an unusually large number of genes annotated with GO terms related to photosynthesis, catalytic activity, kinase activity, cell cycle, and DNA binding (Supplemental Data File S4). The “late high” profile (#6), contained genes with expression patterns complementary to the early high profile, with genes reaching their peak expression in the pink and ripe fruit stages. The genes in this cluster were enriched with GO terms related to transport, sugar metabolism, and catalytic activity, but had a lower than expected proportion of genes annotated to term DNA metabolic process. Only 1% of DNA metabolic process genes were in the late high cluster. These two complementary profiles highlight the diverse biological processes underway in ripe, fully mature fruit versus immature, rapidly growing fruit [55, 57-59]. In the early stages, cell growth and cell division are underway, while in later stages, sugars and other metabolites are being imported into the fruit and cell growth happens largely through cell expansion, not cell division.

The next two most significant clusters also had complementary patterns. The “green high” (#0) cluster included around 1,200 genes that peaked in expression in the mature green stage. This cluster contained unusually many genes annotated with terms related to metabolic processes, transport, DNA binding, and secondary metabolism, suggesting that the green fruit stage represents a developmental transition and also is rich in production of secondary metabolites. The “green low” cluster (#2) contained around 1,500 genes but was significantly enriched with only one term: catalytic activity. Interestingly, genes annotated with this term were unusually prevalent among all genes in all four profiles, suggesting that fruit development is rich in biosynthetic processes. However, the nature of these biosynthetic processes varies among stages, beginning with photosynthetic processes in the early stages and transitioning to secondary metabolism in the later stages. Figures 5D through 5G show RNA-Seq coverage graphs for genes exemplifying the four profiles.

### Transcript variation during fruit development and ripening

As described above, nearly one quarter of multi-exon genes were associated with multiple gene models corresponding to putative products of alternative promoters, alternative splicing, or alternative polyadenylation sites. Any or all three of these mechanisms of alternative transcription can play a role in developmental regulation of gene expression. To test whether alternative transcription was developmentally regulated during berry fruit development and ripening, we used the the CuffDiff program, a companion program to Cufflinks, to identify genes whose patterns of splicing, promoter use, or polyadenylation changed during the development time series. Cuffdiff reported more than 700 genes as undergoing some type of differential transcript variation between pairs of conditions. To assess the results, a subset of the highest confidence CuffDiff results was selected at random and manually inspected using IGB. RNA-Seq read alignments were loaded into the viewer alongside the gene models and read support for alternative transcripts was compared between samples. In most cases involving alternative splice sites, the region that was different between alternative transcripts had small numbers of supporting reads, typically fewer than ten reads per splice junction. By contrast, examples of alternative promoter use (Supplemental Figure 4A) and alternative 3’ ends (Supplemental Figure 4B) had much stronger support. In these instances, typically there were dozens of reads supporting one or both alternatives in each sample. Thus, the CuffDiff analysis was able to identify differential use of alternative promoters and alternative transcription sites but was not as good at distinguishing differential splicing.

We therefore used an alternative approach based on the ArabiTag algorithm [60] to test specifically for differential splicing between stages. For this, a splicing score was calculated that represented the percentage of spliced reads supporting alternative splice site choices from differentially spliced regions. Hierarchical clustering of splicing scores found that ripe and pink berry samples formed a cluster, mature green berries formed a cluster, while the cups and pad stages were intermixed (Figure 6A). Interestingly, the P1 ripe fruit sample was an outlier and formed a distinct cluster apart from the others; this was consistent with previous results in which P1 clustered apart from P2 and P3 in an MDS plot (Figure 5B). Nonetheless, pairwise comparisons of average splicing score found for most alternatively spliced genes, the relative abundance of splice forms was consistent between stages, with some outliers (Figure 6B), and annotated spliced variants were co-expressed. Statistical testing of the splicing score supported this observation, identifying around 90 genes with developmentally regulated differential splicing, including some with predicted functions related to splicing. These included CUFF.35730 (Figure 6C), which was similar to splicing-related transformer-SR ribonucleoproteins from many plant species. The best Arabidopsis match (AT4G35785) is one of two transformer-like genes in Arabidopsis; both genes (AT4G35785 and SR45a) contain alternatively spliced “toxic exons” that introduces a premature stop codon (PTC), and splicing of the toxic exon in SR45a is sensitive to stress [61]. Differential inclusion of a PTC-containing toxic exon appears to be conserved in blueberry, as CUFF.35730 also contained a “toxic exon” that introduced a stop codon. According to the RNA-Seq data, the full-length, exon-skipped form represented a higher percentage of the splice variants in cup and mature green fruit stages, while the exon-included form was less abundant in pink fruit. Thus splicing patterns in blueberry during fruit development and ripening vary by stage, similar to overall gene expression levels.

**Figure 6.**
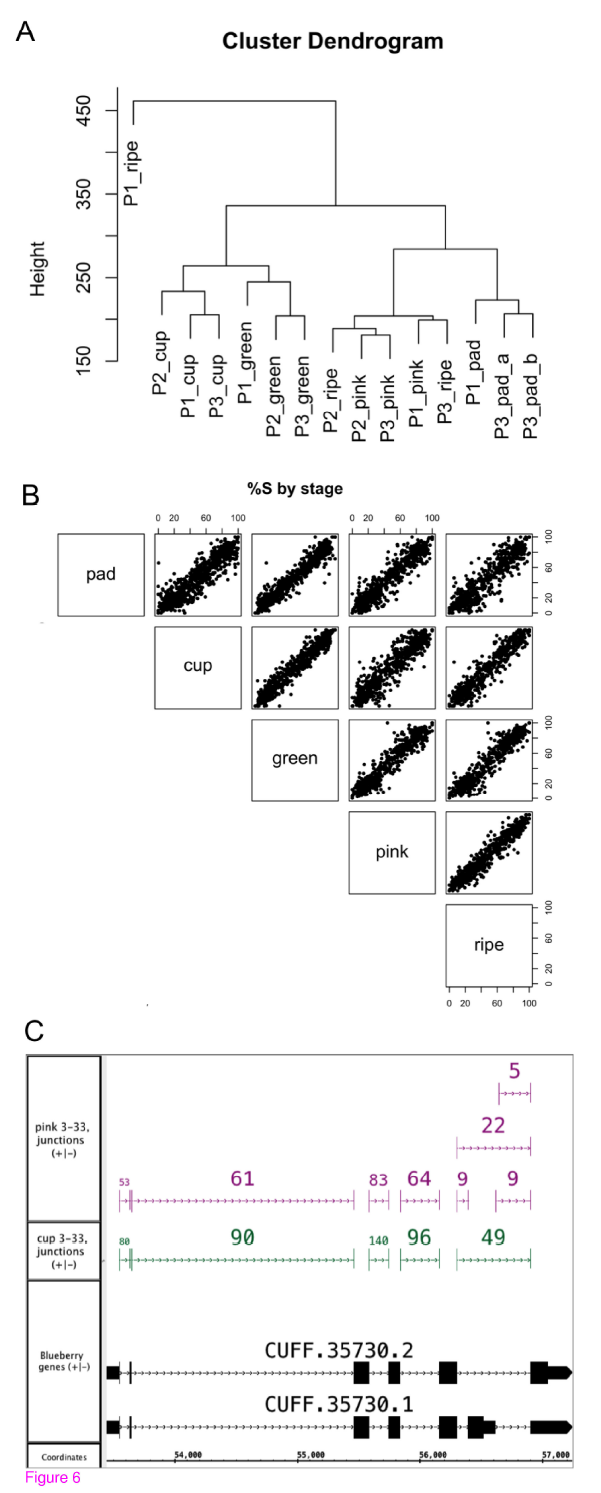
Stage specific alternative splicing. **(A)** Clustering samples by similarity of splicing patterns. (**B**) Scatter plots showing the relationship between average splicing index across different stages. (**C**) Stage-specific alternative splicing in a gene encoding a putative splicing regulator.

### Comparison to prior sequencing studies of blueberry

Prior to now, several studies have been published which used Sanger [17, 62], 454 [16], or Illumina sequencing [63] to characterize blueberry genes and profile gene expression changes. Two studies used *de novo* transcriptome assembly procedures to assemble transcript models, which they then annotated using similar methods to those described here. At the time of writing, only a subset of the data reported in the Illumina-based study was publicly available [63], but the others have made their raw sequence “reads” available as part of dbEST or the Short Read Archive. To provide a resource to researchers and also assess the completeness of the new transcriptome assembly reported here, we aligned publicly available EST, 454 and Illumina sequences onto the blueberry assembly and compared the alignments to the non-redundant set of gene models generated by Cufflinks and the ab initio gene prediction programs. Of the 22,415 Sanger ESTs from dbEST, 19,967 (89%) were aligned onto the draft genome assembly. A comparable percentage of sequences from the much larger 454 sequencing study [16] were also aligned. More than 85% of these 454-based ESTs aligned to the genome, and around 75% of the 454 ESTs overlapped over one or more berry gene models. Similar results were obtained for data from the Illumina-based study [63]. We obtained around 15.5 million read pairs from the Short Read Archive (SRA) using object number SRR942391. Around 55.4% were mapped as proper pairs while another 13.28% were mapped as singletons. Of the mapped reads on the genome, more than 95% overlapped a blueberry gene model. Thus the non-redundant gene models presented here encompass the vast majority of previously generated sequences from blueberry.

The Sanger-EST based study published by Zifkin and co-workers reported in-depth analysis of several genes involved in anthocyanin and proanthocyanidin biosynthesis, two distinct branches of flavonoid biosynthesis [17]. Proanthocyanidins (PAs), also called condensed tannins, are colorless, have a bitter taste, and may play a role in defense against disease and in discouraging consumption of immature berries. Consistent with this, Zifkin et al. found that PA genes were highly expressed in early stages of berry development, while genes specific to anthocyanin biosynthesis were most highly expressed following onset of ripening. Moreover, genes required for synthesis of precursors common to both pathways exhibited a bi-phasic expression profile, where expression was highest in early and late stages and lowest in the intermediate stages. To assess the degree of correspondence between this earlier, EST-based study and the current RNA-Seq based study, EST accessions were obtained from supplementary data [17], their sequences were aligned onto the blueberry genome, and the alignments were used to identify the corresponding blueberry gene models. Alignments were also made available as part of the IGBQuickLoad data repository to enable searching and browsing of EST data. Mappings between ESTs and blueberry gene models are shown in Table IV.

**Table IV.**
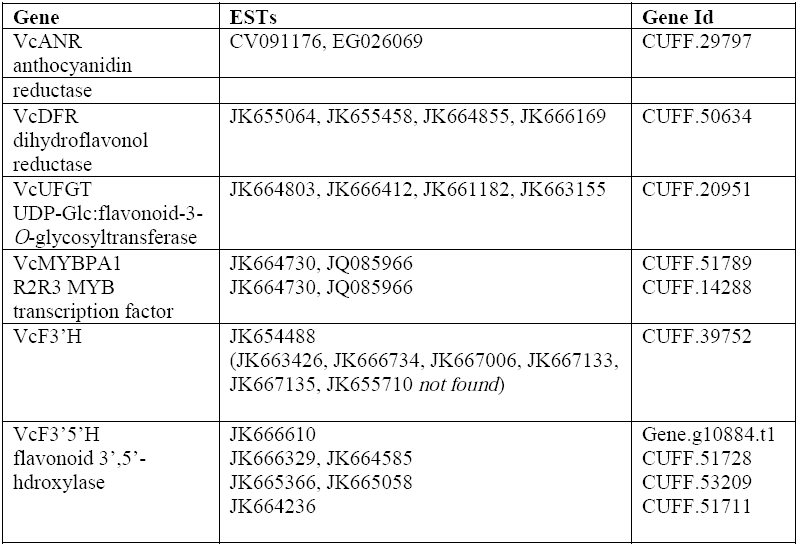
Correspondence between blueberry flavonoid biosynthesis ESTs and blueberry gene models. Previously identified ESTs encoding flavonoid biosynthesis enzymes and regulators were aligned to the blueberry genome. Alignments were visualized alongside the blueberry gene models in Integrated Genome Browser to identify newly annotated genes for each EST.

In general, the RNA-Seq results were consistent with the EST-based study. As shown in Supplemental Figure 5, RNA-Seq based expression patterns for genes encoding VcANR (anthocyanidin reductase), VcDFR (dihydroflavonol reductase), and VcUFGT (UDP-Glc:flavonoid-3-*O*-glycosyltransferase) were similar to those reported previously. Interestingly, the gene corresponding to VcUFGT was CUFF.20951, which was also the most highly expressed gene in the anthocyanin biosynthesis pathway according to the RNA-Seq data (Figure 4A). Zifkin et al also identified ESTs encoding a Myb family transcription factor with biphasic expression and showed it was able to activate an ANR promoter from poplar, demonstrating it is a likely regulator of the PA pathway in blueberry. The EST reported for this gene (JK664730) mapped to two different genes in the blueberry genome assembly, only one of which (CUFF.51789) exhibited the previously reported biphasic expression pattern. The other gene (CUFF.14288) had an expression profile more similar to the “high early, low late” cluster #1 in Figure 5C, suggesting it may co-regulate PA biosynthetic genes in early fruit stages.

## Discussion

In recent years high-throughput sequencing technologies have been used to investigate the transcriptomes of numerous plant species, including many for which no reference genome sequence is available (reviewed in [64]). As sequencing accuracy improves and the cost of sequencing drops, more such projects will become feasible. However, making use of these new data can be challenging when a reference genome is not available to guide the assembly of sequence reads into contigs that are long enough to support functional annotation. Previously, we attempted *de novo* assembly of RNA-Seq reads, but technical difficulties related to the large amount of repetitive sequences in the data stymied these attempts (data not shown). Although the blueberry genome assembly used here is a draft, it was nonetheless complete enough to enable formation of high-quality gene models that could be annotated with GO terms, pathways, and protein homologies. Moreover, the gene models obtained were similar in structure and number to those of many other well-annotated transcriptomes, with one exception, which was that many apparently single-exon genes with no convincing homology to known protein sequences were obtained. Determination of the function of these single-exon, apparently non-protein coding genes is beyond the scope of the current study, but it seems likely that many of these genes may represent long non-coding RNAs, pseudogenes, miRNAs, or other genes whose primary product is RNA and not protein. This possibility does not seem unlikely as other non-coding RNAs as well as pseudogenes have been observed to be expressed in other similarly prepared RNA-Seq libraries [27, 65].

Interestingly, blueberry sequences had highest overall percent identity to sequences from the grape vine *Vitis vinifera*, a basal eudicot with a slow mutation rate. The next most similar species in terms of percent identity was the castor bean *Ricinis communis*, followed by poplar, and then by tomato, the only other asterid species in the RefSeq database at the time this analysis was done. The 454-based transcriptome analysis published by Rowland and co-workers observed an almost identical result [16], as did another group that annotated a *de novo* transcriptome assembly for the tea plant *Camilia sinensis*, another *Ericales* species [66]. These results seem to suggest that blueberry and perhaps even other *Ericales* species are closer to grape than to tomato; however, sequence similarity does not necessarily imply phylogenetic closeness, and a more likely explanation is that grape has diverged at a slower rate than tomato and so is more similar to the last common ancestor of grape and the *Ericacae*. It is also a woody, deciduous plant that makes berry-like fruit and so shares some morphological characteristics with blueberry. The same explanation may also apply to poplar, also a slow-evolving, woody, deciduous plant. However, neither of these explanations serve to explain why castor been, an herbaceous plant with a rapid generation time, appears to be more closely related to blueberry at the sequence level. More recently, phylogenetic analysis of mitochondrial and chloroplast sequences from cranberry (*Vaccinium macrocarpon*) confirmed that cranberry belongs in the asterid grouping and is clearly more closely related to other asterids than grape [67]. It is beyond the scope of this paper to determine correct placement of blueberry relative to these other groups. Instead, the data provided here may provide a means for experts in phylogenetic relationships to investigate these and other questions related to the evolution of flowering plants.

Another important contribution is that for the first time, we provide necessary resources for performing high-throughput gene expression studies in blueberry. The gene models provided here are a key resource for processing and analysis of RNA-Seq expression data, and to maximize their usefulness, we made the RNA-Seq data analysis described here available as part of an open source repository of data files and source code [68]. To complement the gene structures, we provided Gene Ontology and PlantCyc-based enzyme annotations for more than half the protein-coding gene models. The GO and enzyme annotations will provide critical resources for future studies, especially high throughput studies such as RNA-Seq in which statistical analysis typically identifies many hundreds or even thousands of differentially expressed genes. As shown here, being able to identify categories of over- or under-represented in large gene lists enables deeper understanding of biological differences. Moreover, the pathway annotations may further enable interpretation as well as functional prediction. Prior analysis of the AraCyc database together with a compendium of microarray expression data showed that enzymes belonging to the same metabolic pathway are often highly co-expressed, a property that enables identification of missing players in metabolic or regulatory pathways through large-scale analysis of expression data [69, 70]. This approach has been used in many studies that used co-expression to identify candidate genes, leading to a deeper understanding of genes involved in metabolic pathways. Therefore, one important long-term result from the RNA-Seq analysis presented here, along with the pathway predictions, is that we have laid the groundwork for further studies identifying other enzymes in high value pathways. This work may be particularly relevant to anthocyaninin biosynthesis, since different types of anthocyanins have different effects on mammalian biology, possibly due to differences in bioavailability. Another benefit of these data is that because the RNA-Seq data were from an autotetraploid blueberry, aligned onto a diploid genome sequence, it may be possible to use these data to investigate allele-specific gene expression, using SNPs or other polymorphisms to distinguish alleles. This and other studies may become possible thanks to the databases and data sets presented there, and our future work will focus on improving these resources to enable better understanding of metabolic pathways that are active in blueberry.

## Methods

### Analysis code availability

R Markdown files, python scripts, and shell scripts used in data analysis and data processing are version-controlled in the git-based repository at http://www.bitbucket.org/lorainelab/blueberrygenome. The repository contains files documenting which subsections of the repository were used in the analyses described above. Note that we will continue to update and modify the code repository to meet the needs of users; however, readers interested in retrieving older versions of the repository that existed at the time of publication can do so using the git program, which is well documented at http://www.bitbucket.org and many other sites.

### Expression analysis

The number of reads per gene was counted using the samtools view –c command. All single-mapping reads that overlapped a gene region were counted, and gene regions were defined as the smallest start and largest end position of transcripts annotated to the gene. For each comparison, the method “exactTest” with tagwise dispersion from Bioconductor library edgeR [71] was used to identify differentially expressed genes. False discovery rate for each comparison was calculated using the p.adjust method in R, with option “BH” for Benjamini and Hochberg. For clarity, all results reported here applied an FDR cutoff 0.001. Code used to perform differential expression analysis is available in the project bitbucket repository; readers interested in exploring other cutoff options can obtain copies of source code files, edit parameters in the file, and re-run the analysis. Gene Ontology terms that were unusually enriched among differentially expressed genes were identified using GOSeq [72], also from Bioconductor.

### Analysis of alternative splicing with CuffDiff

The non-redundant gene set was used as input to cuffcompare, which produced a file (cuffcmp.combined.gtf) suitable for use as input to cuffdiff. The cuffdiff was then run using the full set of alignments (both multi- and single-mapping) obtained for the blueberry development and ripening time course data set with cuffcomp.combined.gtf file as reference gene models. For analysis of splicing differences, the file splicing.diff output by cuffdiff was used. It was observed that the most significant splicing differences reported by cuffdiff all had the same low p-value (10^−5^) and so these were selected for further analysis. A random subset of the splicing differences from genes that were annotated by the BLASTX step above was selected for visual inspection in IGB.

### Analysis of alternative splicing with ArabiTag

The non-redundant gene set, BAM files with read alignments, and junction files produced by the FindJunctions program (https://bitbucket.org/lorainelab/integrated-genome-browser tools package) were provided to the AltSpliceAnalysis software (https://bitbucket.org/lorainelab/altspliceanalysis), which is based on the ArabiTag algorithm described in [60]. The software identifies annotated alternative splicing events and then uses the RNA-Seq data to count reads supporting alternative splicing choices associated with each event. Data analysis code written in R was used to calculate the percentage of support (%S) for each variant, using gapped reads to support alternative donor or acceptor sites and non-gapped reads to support retention of introns. To identify differential splicing between conditions, a t test was performed on splicing scores. Differential splicing was tabulated and highest confidence differences were manually inspected using Integrated Genome Browser.

## Supplemental Data

The following materials are available in the online version of this article.

**Supplemental Data File S1**. RNA-Seq gene expression data in reads per million kilobase per gene.

**Supplemental Data File S2**. Pathway and enzyme annotations for blueberry genes and transcripts.

**Supplemental Data File S3.** Gene Ontology annotations for blueberry genes and transcripts.

**Supplemental Data File S4**. GO term enrichment analysis of differentially expressed genes.

**Supplemental Data File S5**. GO term enrichment analysis of genes from STEM clusters 0, 1, 2, and 6.

### Acknowledgments

Jim Ballington and the North Carolina Department of Agriculture Piedmont Research Station provided blueberry plants. Lisa J. Rowland donated genomic DNA used in genomic sequencing. Shira Stav and Tyler Estrada helped collect berries used to prepare libraries for RNA sequencing. We thank Nate Watson, Ra’ad Ghabaraih, and Cory Brouwer for contributions to early *de novo* transcriptome assembly efforts not reported here. A grant (to AL) from UNC General Administration provided financial support. Integrated Genome Browser software used in the study was supported by NIH grant 1R01RR032048-01 (to AL). We also express our thanks to the reviewers whose suggestions enabled us to improve the manuscript.

## Author Contributions

AL designed the study, analyzed data, and wrote the manuscript. AE collected berry samples, prepared libraries for sequencing, analyzed data, and helped write the manuscript. IB collected berry samples and analyzed data. KP collected blueberry samples, prepared libraries for sequencing, and analyzed data from the 454 sequencing experiments. VG and SA developed the genome annotation pipeline used to generate blueberry gene models. VG made blueberry gene models, analyzed data, annotated blueberry genes with Gene Ontology terms, and helped write the manuscript. MM analyzed data. RR and AB provided early access to the blueberry genome sequence. All authors read and helped edit the manuscript.

## Additional files provided with this submission

Additional file 1: berry_RPKM.tsv.gz, 6737K http://www.gigasciencejournal.com/imedia/1759828185140931/supp1.gz

Additional file 2: locus2pathway_60_45.tsv.gz, 61K http://www.gigasciencejournal.com/imedia/7414054511409310/supp2.gz

Additional file 3: S4-GO.tsv, 1566K http://www.gigasciencejournal.com/imedia/9154200061409310/supp3.tsv

Additional file 4: SFigures.pdf, 3108K http://www.gigasciencejournal.com/imedia/5063485221409310/supp4.pdf

Additional file 5: GO-DE.tsv.gz, 8K http://www.gigasciencejournal.com/imedia/7048455931515400/supp5.gz

Additional file 6: S5-GO-Cluster.tsv.gz, 1K http://www.gigasciencejournal.com/imedia/1338250947151540/supp6.gz

Additional file 7: BlueberryTranscriptome-TrackedChanges.docx, 209K http://www.gigasciencejournal.com/imedia/1516791598156311/supp7.docx

